# Chronic activation of fear engrams induces extinction-like behavior in ethanol-exposed mice

**DOI:** 10.1101/2020.07.01.182782

**Authors:** Christine Cincotta, Nathen J. Murawski, Stephanie L. Grella, Olivia McKissick, Emily Doucette, Steve Ramirez

## Abstract

Alcohol withdrawal directly impacts the brain’s stress and memory systems, which may underlie individual susceptibility to persistent drug and alcohol-seeking behaviors. Numerous studies demonstrate that forced alcohol abstinence, which may lead to withdrawal, can impair fear-related memory processes in rodents such as extinction learning, however the underlying neural circuits mediating these impairments remain elusive. Here, we tested an optogenetic strategy aimed at mitigating fear extinction impairments in male c57BL/6 mice following exposure to alcohol (i.e., ethanol) and forced abstinence. In the first experiment, extensive behavioral extinction training in a fear-conditioned context was impaired in ethanol-exposed mice compared to controls. In the second experiment, neuronal ensembles processing a contextual fear memory in the dorsal hippocampus were tagged and optogenetically reactivated repeatedly in a distinct context in ethanol-exposed and control mice. Chronic activation of these cells resulted in a context-specific, extinction-like reduction in fear responses in both control and ethanol-exposed mice. These findings suggest that while ethanol can impair fear extinction learning, optogenetic manipulation of a fear engram is sufficient to induce an extinction-like reduction in fear responses.

---

More than 50 percent of people who undergo formal treatment for an alcohol use disorder (AUD) relapse before they can reach a year of sobriety (Connor et al. 2016). A major factor contributing to the susceptibility of relapse involves the direct, negative impact that withdrawal from alcohol has on stress and memory systems (Koob, 2008; Staples & Mandyam, 2016). Exposure to drug-associated cues, including contextual cues, can trigger memories of past alcohol-drinking experiences, increase levels of stress-related hormones, and potentiate relapse-related behavior (Chaudhri et al. 2008). The hippocampus, a key brain region involved in both stress and memory processes, is critical in the formation of drug-context associations and may therefore be a potential neural target to mitigate stress- and memory-induced relapse behavior in AUDs (Goode & Maren, 2019).

Exposure-based therapies – founded upon the behavioral principles of extinction learning (Goode and Maren, 2019; Bouton, 2019; Clem and Schiller, 2016) – have been somewhat successful in the clinical treatment of anxiety, stress, and addiction disorders (Kaplan, et al. 2011; Mellentin, et al. 2017). In extinction learning, exposure to a conditioned stimulus (context) in the absence of an unconditioned stimulus (shock) leads to a reduction in conditioned behavior (e.g. freezing in the case of fear-conditioning) (Bouton, 2019). Although exposure-based therapies, which operate on the principles of extinction learning, have been used as a treatment option for addiction, their moderate success is likely related to the fact that preclinical studies demonstrate significant impairments in extinction processes in rodent models of alcohol (i.e., ethanol) use and withdrawal. For example, ethanol dependence can induce increased resistance to extinction where more sessions to reach an extinction criterion are required following intermittent ethanol exposure in adult (Gass et al. 2017) and adolescent rats (Gass et al., 2014). Furthermore, rodent models also demonstrate that ethanol-related cues as well as stress can induce reinstatement and context-dependent renewal of ethanol-associated lever pressing following extinction (Burattini et al. 2006; Chaudhri et al., 2008; Keistler et al. 2017).

These data suggest that aspects of ethanol withdrawal can promote ethanol-seeking behavior potentially through mechanism of enhanced stress (Koob, 2008) where disruption of extinction occurs, thereby potentially hampering clinical attempts to utilize extinction processes to treat AUDs.

In rodents withdrawn from ethanol, preclinical studies also show impairments in memory processes unrelated to the ethanol itself (e.g. fear learning). These deficits include impaired extinction learning, heightened, stress-induced reinstatement, and increased fear generalization (Holmes, et al. 2012; Bertotto, et al. 2006; Broadwater & Spear, 2013; Scarlata et al. 2019). Further, fear memories formed prior to ethanol exposure and withdrawal also fail to extinguish (Quinones-Laracuente, et al. 2015), suggesting that withdrawal from ethanol augments the retrieval of fear memories and disrupts general mechanisms of extinction learning. If extinction impairments in ethanol-withdrawn rodents reflect retrieval deficits related to the extinction memory (e.g., enhanced fear generalization) this may be manifested as increased freezing in a neutral context (Jasnow, et al. 2017). In this case, targeting retrieval processes may prove effective at mitigating the extinction impairments observed following withdrawal from ethanol.

Here, we employ a strategy that drives context-specific associative learning that may overcome these retrieval errors by targeting the original fear memory (Chen et al, 2019). Fear memory retrieval is a dynamic process that can be measured at the neuronal, circuit, and behavioral level. Moreover, a behavioral strategy to suppress fear and induce extinction learning involves returning a rodent to the fear-conditioning context in the absence of shock (Goode & Maren, 2019). Repeated sessions where retrieval of a fear memory in the absence of the unconditioned stimulus leads to an overall reduction in conditioned response (i.e., freezing (Goode & Maren, 2019). An alternative method involves an artificial strategy, which attempts to optogenetically activate a fear engram – the neuronal ensemble active during acquisition which undergoes plasticity, and which reactivation of facilitate retrieval (Liu, et al. 2012; Ramirez, et al. 2015; Chen, et al. 2019; Josselyn and Tonegawa 2020). Promisingly, optogenetic activation of cells in the dorsal dentate gyrus (dDG) subregion of the hippocampus that were previously active during fear learning is sufficient to activate the neuronal and behavioral expression of memory recall (Liu, et al. 2012; Ramirez, et al. 2013; Ramirez, et al. 2015). Recently, we demonstrated that repeated (or “chronic”) optical activation of a fear memory in the dDG leads to a context-specific, extinction-like reduction in freezing (Chen, et al., 2019). Therefore, we asked if repeated activation of a hippocampal fear engram could mitigate forced abstinence-induced fear extinction deficits, possibly caused by acute withdrawal, in a context-specific manner.

We first utilized a behavioral strategy to evaluate aberrant fear extinction in a mouse model of chronic ethanol exposure and forced abstinence. All subjects were treated in accordance with protocol 17-008 approved by the Institutional Animal Care and Use Committee at Boston University. Adult male mice received intraperitoneal (IP) injections of 30% ethanol (vol/vol) in saline (0.9%) at a dose of 1.5 g/kg [EtOH] or saline [Sal] for 5 consecutive days, followed by a two-day forced abstinence period (Quinones-Laracuente et al., 2015, Pina and Cunningham, 2016). Twenty-eight male C57Bl6/J mice (Sal=14, EtOH=14) were fear conditioned in two distinct contexts, Context A (Ctx A) and Context B (Ctx B), extinguished in Ctx A, and tested in both Ctx A and B as previously described (Chen, et al. 2019; Figure 1A).

**Figure 1:**
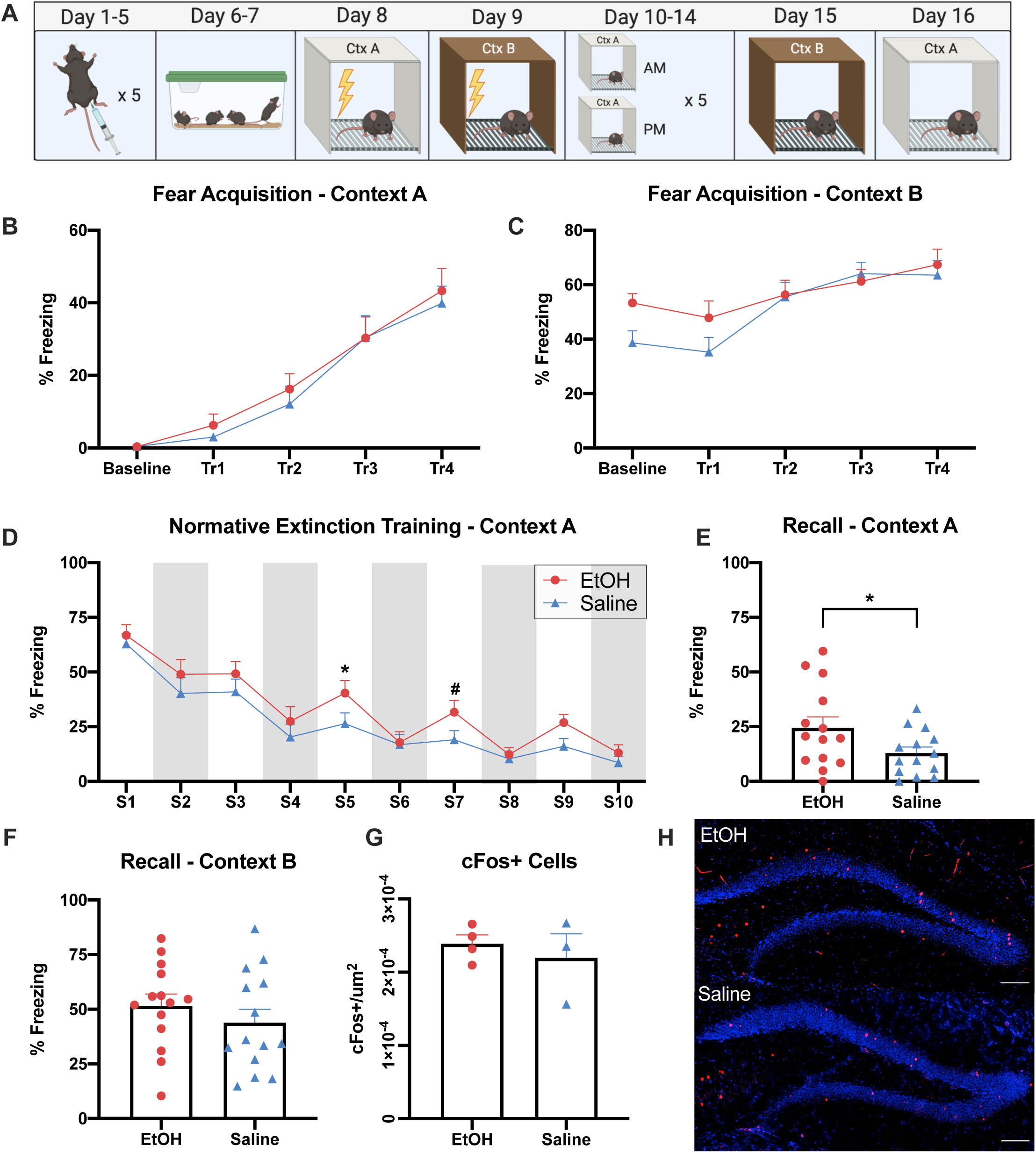
Ethanol exposure and forced abstinence impairs normative extinction learning. A) Behavioral design for ethanol (EtOH) or saline exposure, forced abstinence, fear conditioning, extinction, and recall. B) Fear acquisition of EtOH-exposed and Sal-control mice in Ctx A. No differences between groups (two-way RM ANOVA, trial) (F _(4,72)_ = 54.98, p <0.0001). C) Fear acquisition of EtOH-exposed and Sal-control mice in Ctx B. No differences between groups (two-way RM ANOVA, trial) (F _(4,72)_ = 15.54, p <0.0001). D) Normative extinction training revealed an impaired ability to attenuate a fear memory in EtOH-exposed mice (two-way RM ANOVA, session: F _(9,234)_ = 65.47, p <0.0001; S5 EtOH *vs*. Sal: Tukey’s HSD, *p = 0.049; S7 EtOH *vs*. Sal: Tukey’s HSD, #p = 0.082). E) Recall in context A following extinction reveals increased freezing in EtOH-withdrawn mice (unpaired t-test, one-tailed, t _(26)_ = 2.020, p = 0.0269). F) In contrast, no group differences were found in freezing during the recall test in context B (unpaired t-test, two-tailed, t _(26)_ = 0.968, p = 0.3418). G) Quantification of dDG cFos+ cells during recall in context A reveals no significant difference in activation (unpaired t-test, two-tailed, t _(5)_ = 0.629, p = 0.5567). H) Representative histology for cFos+ quantification. Scale bar = 100 um.

We observed no group differences in fear acquisition in Ctx A (Figure 1B) however, there was a main effect of trial demonstrating increases in freezing in the latter trials compared to baseline and earlier trials (two-way RM ANOVA; F _(4,72)_ = 54.98, p <0.0001). This was also the case in Ctx B (Figure 1C) (two-way RM ANOVA; F _(4,72)_ = 15.54, p <0.0001). Following fear acquisition, mice underwent 10-min extinction sessions in Ctx A (2 sessions per day for 5 days). Across extinction, we observed a gradual decrease in freezing levels (Figure 1D) (two-way RM ANOVA, session) (F _(9,234)_ = 65.47, p <0.0001). Given our effects were stronger in the morning extinction sessions, we followed up this analysis by looking at these sessions specifically and found a similar effect (two-way RM ANOVA, session) (F _(4,104)_ = 84.65, p <0.0001). Post-hoc analyses revealed that EtOH-exposed mice were impaired in extinction learning. On the third day (S5), EtOH mice froze significantly more than Sal mice (Tukey’s HSD, p = 0.049). As the extinction trials progressed these mice eventually started to reach normative levels (S7: Tukey’s HSD, p = 0.082) (Figure 1D).

Following extinction, mice were tested for contextual fear memory recall in Ctx A (Figure 1E). As expected, freezing levels of EtOH-exposed mice were significantly higher relative to Sal-control mice (unpaired t-test, one-tailed, t _(26)_ = 2.020, p = 0.0269). In contrast, no group differences were found in freezing during the recall test in Ctx B (unpaired t-test, two-tailed, t _(26)_ = 0.968, p = 0.3418) (Figure 1F). Following behavior, cFos+ cells in the dDG were quantified as a proxy of activation during recall in Ctx A (Figure 1G-H). We observed no significant difference in this cellular marker of activity during recall (unpaired t-test, two-tailed, t _(5)_ = 0.629, p = 0.5567) (Figure 1G). Together, these data show that EtOH-exposed mice take longer to extinguish a fear-conditioned freezing response, and a higher degree of freezing during recall which was context-specific despite comparable levels of cFos activation in the dDG.

Our previous work showed that chronic stimulation of cells in the dorsal hippocampus active during the formation a fear memory led to an extinction-like, context-specific reduction in fear behavior in mice (Chen et al. 2019). Therefore, we asked if chronic optogenetic activation of a fear engram in the dDG could facilitate extinction learning in EtOH-exposed mice. Forty-four adult male mice were placed on a diet containing doxycycline (40 mg/kg; Dox). Briefly, mice were injected with a virus cocktail [pAAV_9_-c-Fos-tTA and either pAAV_9_-TRE-ChR2-eYFP (ChR2) or pAAV_9_-TRE-eYFP (eYFP)] into the dDG followed by bilateral optic fiber implantation (Figure 2A-B; see Ramirez et al., 2015; Chen et al. 2019).

**Figure 2:**
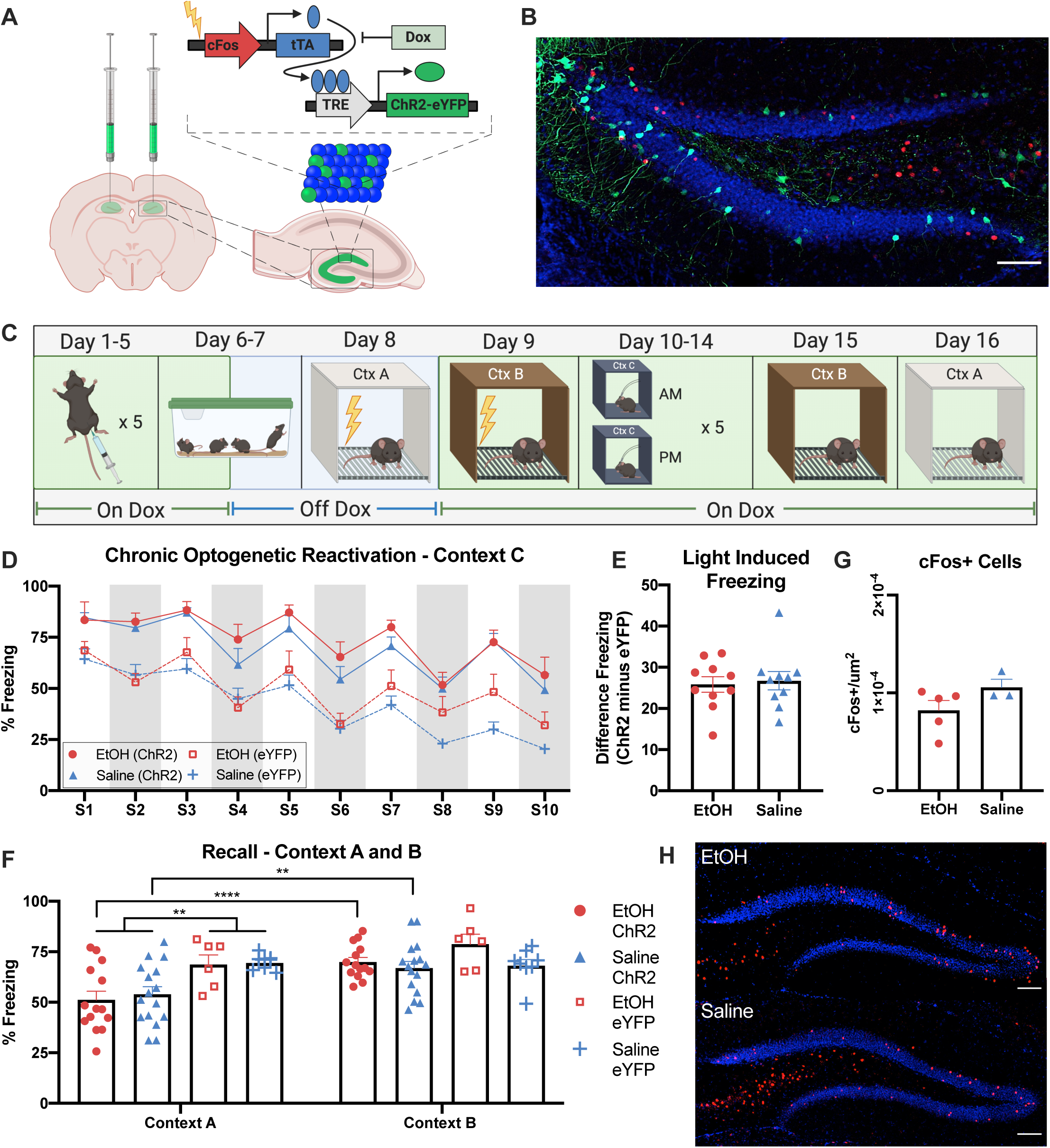
Chronic optogenetic activation reduces freezing in ethanol-exposed mice. A) Schematic of viral strategy. A viral cocktail of AAV9-c-Fos-tTA and either AAV9-TRE-ChR2-eYFP or AAV9-TRE-eYFP was infused into the dDG for activity-dependent transcription of ChR2 or eYFP. B) Representative histology for activity-dependent tagging of contextual fear engrams. Scale bar = 100 um. C) Behavioral design for ethanol (EtOH) or saline exposure, forced abstinence, fear conditioning, chronic optogenetic activation, and recall. D) Freezing levels during chronic activation of a contextual fear engram in neutral context C. While there is an overall decrease in freezing across all sessions, we found no significant group differences based on treatment for both ChR2 and eYFP groups (three-way RM ANOVA) (session F_(9, 216)_ = 44.57, p<0.0001, virus F_(1, 216)_ = 38.74 p<0.0001). E) Difference score for light-induced freezing. For each optogenetic activation session, the difference in freezing between ChR2 and eYFP groups for both EtOH and Sal treatments were calculated. Average difference in freezing is equal for both treatment groups (unpaired t-test on difference score, two-tailed, t _(18)_ = 0.3030, p = 0.7654). F) Recall in context A (Ctx A) reveals a significant decrease in freezing levels of ChR2 EtOH and saline mice relative to eYFP control mice. Significant context by treatment interaction (three-way RM ANOVA) (F _(1,40)_ = 4.274, p = 0.0452) (Tukey’s HSD, EtOH-ChR2: p < 0.001; Sal-ChR2: p = 0.032). Additionally, this decrease in freezing was context specific, with significantly lower freezing levels of EtOH-ChR2 and Sal-ChR2 mice in Ctx A relative to Ctx B. Significant context x virus interaction (three-way RM ANOVA) (F _(1,40)_ = 7.721, p = 0.0083) (Tukey’s HSD: Ctx A EtOH-ChR2 *vs*. Sal-ChR2 p < 0.001) suggesting that chronic optogenetic activation induced a context-specific decrease in freezing. G) Quantification of dDG cFos+ cells during recall in Ctx A reveals no significant difference in activation (unpaired t-test, two-tailed, t _(6)_ = 1.6, p = 0.1606). H) Representative histology for cFos+ quantification. Scale bar = 100 um.

Following recovery, mice received 5 days of ethanol or saline treatment. As in the previous experiment, mice underwent a two-day forced abstinence period prior to fear conditioning. Mice were taken off-Dox 24 hours prior to fear conditioning in Ctx A, and immediately put back on-Dox after tagging for subsequent fear conditioning in Ctx B. Over the next five days, mice were placed into a distinct context (Ctx C) and received repeated light-stimulation (473nm, 20 Hz) over a 10-min session, twice a day for five days, as previously reported (Ramirez et al. 2015; Chen et al. 2019). Finally, mice were tested for contextual fear memory recall in both Ctx A and Ctx B (Figure 2C). Analysis of freezing during optogenetic activation sessions revealed a main effect of session, a main effect of virus, and no main effect of treatment (three-way RM ANOVA) (session F_(9,216)_ = 44.57, p<0.0001, virus F_(1,216)_ = 38.74 p<0.0001) (Figure 2D). Differences in freezing between EtOH ChR2 and eYFP groups as well as Saline ChR2 and eYFP groups indicated significant light-induced freezing from reactivation of the initial tagged fear conditioning session (Figure 2D), and equal levels of light-induced freezing were observed in each ChR2 group (unpaired t-test on difference score, two-tailed, t _(18)_ =0.3030, p=0.7654; Figure 2E).

We measured freezing in Ctx A and Ctx B for ChR2 EtOH-exposed and Sal-control mice as well as eYFP controls. Overall, we found a main effect of context, a main effect of virus, and no main effect of treatment. We found a significant context by treatment interaction (three-way RM ANOVA) (F _(1,40)_ = 4.274, p = 0.0452). This effect is driven by the observation that across treatment, there is a significantly lower degree of freezing in Ctx A for ChR2 animals (Tukey’s HSD, EtOH-ChR2: p < 0.001; Sal-ChR2: p = 0.032), suggesting that chronic optogenetic activation induced a context-specific decrease in freezing. This effect was not observed in eYFP mice. Additionally, there was a significant context x virus interaction (three-way RM ANOVA) (F _(1,40)_ = 7.721, p = 0.0083) and this effect was due to the difference in freezing within Ctx A, between the EtOH ChR2 group and the Sal eYFP group (Tukey’s HSD, p < 0.001) and between ChR2 and eYFP mice overall (Tukey’s HSD, ChR2 *vs*. eYFP: p < 0.001) (Figure 2F).

Following behavior, cFos+ cells in the dDG were quantified as a proxy of activation during recall in Ctx A (Figure 2G-H). We observed no significant difference in cellular activity during recall (unpaired t-test, two-tailed, t _(6)_ = 1.6, p = 0.1606) (Figure 2F). Together, these data demonstrate that chronic activation of a tagged dDG fear memory leads to a context-specific reduction in freezing in both ChR2 EtOH and Sal mice.

We utilized a behavioral and optogenetic approach to respectively evaluate and reduce aberrant fear extinction responses in mice that experienced ethanol exposure and forced abstinence. Behaviorally, repeated exposure to a fear-associated context led to fear retrieval. Although both groups of mice acquired fear to a similar extent to both contexts, EtOH-exposed mice demonstrated a slower rate of context extinction and an overall moderate impairment in extinction memory relative to Sal-control mice. Next, chronic optogenetic activation of tagged dDG cells induced a context-specific, extinction-like reduction of freezing behavior in both Sal-control and EtOH-exposed mice, but not in eYFP controls. Moreover, freezing levels in the tagged context did not differ between ChR2 EtOH and Sal mice. These data lend credence to the idea that artificially reactivating a fear memory over multiple sessions, is sufficient to reduce fear memories in an extinction-like manner in both Sal-control and EtOH-exposed mice, and points to dDG-mediated engrams as a key node, sufficient to alleviate maladaptive behavioral responses.

We hypothesize that forced abstinence of ethanol may have caused withdrawal-induced changes to stress systems in our mice, which led to a general enhancement in fear learning. It is possible that this increased fear learning was therefore more resistant to behavioral extinction, but susceptible to optogenetic perturbations. Fear extinction impairments in the EtOH-exposed mice reported here dovetails with previous reports of ethanol-withdrawal memory impairments, including heightened fear responses, increased context generalization, impaired extinction, and stress/cue-induced reinstatement in rodents (Holmes, et al. 2012; Bertotto, et al. 2006; Broadwater & Spear, 2013; Quinones-Laracuente, et al. 2015).

Additionally, we found no difference in cFos+ cell quantifications between EtOH-exposed and Sal-control groups in both the normative and optogenetic experiments. Though future studies can increase sample sizes to potentially detect subtle differences across groups, it is likely an upstream brain region mediates the behavioral impairment observed in normative EtOH-exposed mice. For instance, previous studies regarding fear memory retrieval altered by ethanol have found increased cFos in the prelimbic cortex, paraventricular thalamus, and medial central amygdala, and have found these regions function as a circuit during fear retrieval (Quinones-Laracuente et al 2015; Do-Monte et al. 2015). In our experiments, we speculate that by perturbating an upstream hippocampal node implicated in processing the contextual components of a fear memory engram, we optogenetically engaged independent circuitry capable of attenuating fear responses and thus artificially facilitated extinction-like behaviors in both EtOH-exposed and Sal-control mice. In line with this view, and despite our negative cFos data in Figure 2, we propose that our chronic stimulation protocol bypassed numerous brain-wide systems previously implicated in mediating addiction-related behaviors, such as the insular cortex, which governs interoceptive feelings of drug craving and withdrawal (Naqvi & Bechara 2009). Measuring immediate early genes in the insula may reveal differential activity levels between EtOH-exposed and Sal-control mice, which potentially underlie the impaired ability of EtOH-exposed mice to mitigate fear memories.

Moreover, recent studies have shown that discrete populations of cells contributing to a memory engram have distinct cellular activity, synaptic properties, and behavioral results dependent on the immediate early gene used to identify a given cellular ensemble (Sun et al. 2020). Specifically, cFos-mediated engrams have been implicated in promoting memory generalization, such that acute chemogenetic activation of a contextual fear memory led to reduced discrimination between distinct contexts (Sun et al. 2020). In parallel, chronic activation of a fear memory in a neutral context may have promoted generalization between the neutral and conditioned context, as evident by decreased freezing in the conditioned context. Previously reported pharmacological and optogenetic approaches that mitigate stress- and addiction-related behavioral states have demonstrated enduring changes in behavior following sustained, but not acute, activation of neural systems (Friedman, et al. 2014; Ramirez, et al. 2015); likewise, we posit that our chronic activation protocol may enduringly reprogram and/or modify existing engrams and circuits sufficient to alleviate addiction-related behavioral states. Alternatively, chronic optogenetic activation may cause an artificial extinction-like engram to emerge in the DG. Previous studies have demonstrated that extinction recruits a new population of DG cells that compete with the original fear engram to drive each corresponding behavior (Khalaf et al. 2018; Lacagnina et al. 2019; Chen et al. 2019). In our recent study, we observed that chronic stimulation of DG-mediated engrams led to a reduction of cFos activity in the originally tagged population of cells while the DG nonetheless maintained a similar level of overall cFos expression levels in non-tagged populations, suggesting too that a distinct engram had simultaneously emerged—this artificial engram may function to suppress freezing behavior elicited by the original fear engram (Chen et al. 2019, Lacagnina et al 2019).

Taken together, the emergence of novel strategies sufficient to mitigate fear responses holds promising value in better understanding the underlying mechanisms of learning and memory. In particular, modulating addiction-related engrams permits a brain-wide cataloguing of the maladaptive structural and functional changes while pointing to key cellular mechanisms that may be sites of future intervention (Whitaker & Hope 2018). By monitoring and manipulating the cellular, circuit, and systems-wide changes that occur when a brain transitions into a state of drug or alcohol-dependence, we believe it may be possible to artificially and enduringly restore healthy neuronal functioning and corresponding behavioral outputs.

## Acknowledgements

We thank Dr. Joshua Sanes and his lab at the Center for Brain Science, Harvard University, for providing laboratory space within which the initial experiments were conducted, the Center for Brain Science Neuroengineering core for providing technical support, and the Society of Fellows at Harvard University for their support. We also thank Dr. Susumu Tonegawa and his lab for providing the activity-dependent virus cocktail. This work was supported by an NIH Early Independence Award (DP5 OD023106-01), an NIH Transformative R01 Award, a Young Investigator Grant from the Brain and Behavior Research Foundation, a Ludwig Family Foundation grant, the McKnight Foundation Memory and Cognitive Disorders award, and the Center for Systems Neuroscience and Neurophotonics Center at Boston University.

## References

Bertotto, M. E., Bustos, S. G., Molina, V. A., & Martijena, I. D. (2006). Influence of ethanol withdrawal on fear memory: Effect of D-cycloserine. Neuroscience, 142(4), 979–990. doi: 10.1016/j.neuroscience.2006.07.013

Bouton, M.E. (2019). Extinction of instrumental (operant) learning: interference, varieties of context, and mechanisms of contextual control. Psychopharmacology, 236, 7–19.

Broadwater, M., & Spear, L. P. (2013). Consequences of ethanol exposure on cued and contextual fear conditioning and extinction differ depending on timing of exposure during adolescence or adulthood. Behav Brain Res, 256, 10–19. doi: 10.1016/j.bbr.2013.08.013

Burattini, C., Gill, T.M., Aicardi, G., Janak, P.H. (2006). The ethanol self-administration context as a reinstatement cue: acute effects of naltrexone. Neuroscience, 139(3), 877–887.

Chaudhri, N., Sahuque, L. L., & Janak, P. H. (2008). Context-induced relapse of conditioned behavioral responding to ethanol cues in rats. Biol Psychiatry, 64(3), 203–210. doi: 10.1016/j.biopsych.2008.03.007

Chen B., Murawski, N.J., Hamidi, A.B., Merfeld, E., Doucette, E., Grella, S.L., Shpokayte, M., Zaki, Y., Cincotta, C., Fortin, A., Ramirez, S. (2019). Artificially enhancing and suppressing hippocampus-mediated memories. Current Biology, 29, 1–10.

Clem RL, Schiller D. New Learning and Unlearning: Strangers or Accomplices in Threat Memory Attenuation? Trends Neurosci. 2016;39(5):340–351. doi: 10.1016/j.tins.2016.03.003

Connor, J.P., Haber, P.S., Hall, W.D. (2016). Alcohol use disorders. Lancet, 387(10022), 988–998.

Do-Monte, F. H., Quiñones-Laracuente, K., & Quirk, G. J. (2015). A temporal shift in the circuits mediating retrieval of fear memory. Nature, 519(7544), 460–463. https://doi.org/10.1038/nature14030

Gass, J.T., McGonigal, J.T., Chandler, L.J. (2017). Deficits in the extinction of ethanol-seeking behavior following chronic intermitten ethanol exposure are attenuated with positive allosteric modulation of mGlu5. Neuropharmacology, 113, 198–205.

Gass, J. T., Glen, W. B., Jr, McGonigal, J. T., Trantham-Davidson, H., Lopez, M. F., Randall, P. K., Yaxley, R., Floresco, S. B., & Chandler, L. J. (2014). Adolescent alcohol exposure reduces behavioral flexibility, promotes disinhibition, and increases resistance to extinction of ethanol self-administration in adulthood. Neuropsychopharmacology, 39(11), 2570–2583. https://doi.org/10.1038/npp.2014.109

Good, T.D., Maren, S. (2019). Common neurocircuitry mediating drug and fear relapse in preclinical models. Psychopharmacology, 236(1), 415–437. doi: 10.1007/s00213-018-5024-3.

Holmes, A., Fitzgerald, P. J., MacPherson, K. P., DeBrouse, L., Colacicco, G., Flynn, S. M., … Camp, M. (2012). Chronic alcohol remodels prefrontal neurons and disrupts NMDAR-mediated fear extinction encoding. Nat Neurosci, 15(10), 1359–1361. doi: 10.1038/nn.3204

Jasnow, A. M., Ehrlich, D. E., Choi, D. C., Dabrowska, J., Bowers, M. E., McCullough, K. M., … Ressler, K. J. (2013). Thy1-expressing neurons in the basolateral amygdala may mediate fear inhibition. J Neurosci, 33(25), 10396–10404. doi: 10.1523/JNEUROSCI.5539-12.2013

Kaplan, G.B., Heinrichs, S.C., Carey, R.J. (2011). Threatment of addiction and anxiety using extinction approaches: neural mechanisms and their treatment implications. Pharmacology, Biochemistry, and Behavior, 97(3), 619–25.

Keistler, C.R., Hammarlund, E., Barker, J.M., Bond, C.W., DiLeone, R.J., Pittenger, C., Taylor, J.R. (2017). Regulatio nof alcohol extinction and cue-induced reinstatment by specific projections among medial prefrontal cortex, nucleus accumbens, and basolateral amygdala. Journal of Neuroscience, 37(17), 4462–4471.

Khalaf, O., Resch, S., Dixsaut, L., Gorden, V., Glauser, L., & Gräff, J. (2018). Reactivation of recall-induced neurons contributes to remote fear memory attenuation. Science, 360(6394), 1239–1242. https://doi.org/10.1126/science.aas9875

Koob, G. F. (2008). A role for brain stress systems in addiction. Neuron, 59(1), 11–34. doi: 10.1016/j.neuron.2008.06.012

Lacagnina, A.F., Brockway, E.T., Crovetti, C.R., Shue, F., McCarty, M.J., Sattler, K.P., Lim, S.C., Santos, S.L., Denny, C.A., & Drew, M.R. (2019) Distinct hippocampal engrams control extinction and relapse of fear memory. Nat Neurosci 22, 753–761. https://doi.org/10.1038/s41593-019-0361-z

Liu, X., Ramirez, S., Pang, P. T., Puryear, C. B., Govindarajan, A., Deisseroth, K., & Tonegawa, S. (2012). Optogenetic stimulation of a hippocampal engram activates fear memory recall. Nature, 484(7394), 381–385. doi: 10.1038/nature11028

Mellentin, A.I., Skot, L., Neilsen, B., Schippers, G.M., Nielsen, A.S., Stenager, E., Juhl, C. (2017). Cue exposure therapy for the treatment of alcohol use disorders: A meta-analytic review. Clinical Psychology Review, 57, 195–207.

Pina, M.M., Cunningham, C.L. (2017). Ethanol-seeking behavior is expressed directly thorugh an extended amygdala to midbrain neural circuit. Neurobiology of Learning and Memory, 137, 83–91.

Quinones-Laracuente, K., Hernandez-Rodriguez, M. Y., Bravo-Rivera, C., Melendez, R. I., & Quirk, G. J. (2015). The effect of repeated exposure to ethanol on pre-existing fear memories in rats. Psychopharmacology (Berl), 232(19), 3615–3622. doi: 10.1007/s00213-015-4016-9

Ramirez, S., Liu, X., Lin, P. A., Suh, J., Pignatelli, M., Redondo, R. L., … Tonegawa, S. (2013). Creating a false memory in the hippocampus. Science, 341(6144), 387–391. doi: 10.1126/science.1239073

Ramirez, S., Liu, X., MacDonald, C. J., Moffa, A., Zhou, J., Redondo, R. L., & Tonegawa, S. (2015). Activating positive memory engrams suppresses depression-like behaviour. Nature, 522(7556), 335–339. doi: 10.1038/nature14514

Scarlata, M.J., Lee, S.H., Lee, D., Kandigian, S.E., Hiller, A.J., Dishart, J.G., Mintz, G.E., Wang, Z., Coste, G.I., Mousley, A.L., Soler, I., Lawson, K., Ng, A.J., Bezek, J.L., & Bergstrom, H.C. (2019). Chemogenetic stimulation of the infralimbic cortex reverses alcohol-induced fear memory overgeneralization. Scientific Reports 9, 6730. https://doi.org/10.1038/s41598-019-43159-w

Staples, M. C., & Mandyam, C. D. (2016). Thinking after Drinking: Impaired Hippocampal-Dependent Cognition in Human Alcoholics and Animal Models of Alcohol Dependence. Front Psychiatry, 7, 162. doi: 10.3389/fpsyt.2016.00162

Sun, X., Bernstein, M. J., Meng, M., Rao, S., Sørensen, A. T., Yao, L., Zhang, X., Anikeeva, P. O., & Lin, Y. (2020). Functionally Distinct Neuronal Ensembles within the Memory Engram. Cell, 181(2), 410–423.e17. https://doi.org/10.1016/j.cell.2020.02.055

Whitaker, L. R., & Hope, B. T. (2018). Chasing the addicted engram: identifying functional alterations in Fos-expressing neuronal ensembles that mediate drug-related learned behavior. Learning & Memory), 25(9), 455–460. https://doi.org/10.1101/lm.046698.117

